# Contrasting processing tomato cultivars unlink yield and pollen viability under heat stress

**DOI:** 10.1101/2021.04.16.439802

**Authors:** Golan Miller, Avital Beery, Prashant Kumar Singh, Fengde Wang, Rotem Zelingher, Etel Motenko, Michal Lieberman-Lazarovich

## Abstract

The occurring climate change is causing temperature increment in crop production areas worldwide, generating conditions of heat stress that negatively affect crop productivity. Tomato (*Solanum lycopersicum*), a major vegetable crop, is highly susceptible to conditions of heat stress. When tomato plants are exposed to ambient day/night temperatures that exceed 32°C/20°C respectively during the reproductive phase, fruit set and fruit weight are reduced, leading to a significant decrease in yield. Processing tomato cultivars are cultivated in open fields, where environmental conditions are not controlled, therefore plants are exposed to multiple abiotic stresses, including heat stress. Understanding the physiological response of modern processing tomato cultivars to heat stress may facilitate the development of thermotolerant cultivars. Here, we compared two tomato processing cultivars, H4107 and H9780, that we found to be constantly differing in yield performance. Using field and temperature-controlled greenhouse experiments, we show that the observed difference in yield is attributed to the occurrence of heat stress conditions. In addition, fruit-set and seed production were significantly improved in the thermotolerant cultivar H4107, compared with H9780. Despite the general acceptance of pollen viability as a measure of thermotolerance, there was no difference in the percentage of viable pollen between H4107 and H9780 under either of the conditions tested. Therefore, processing tomato cultivars may present a particular case, in which other factors are central for heat stress tolerance. Our results also demonstrate the value of combining controlled with uncontrolled experimental settings, in order to identify heat stress related responses and facilitate the development of thermotolerant processing tomato cultivars.

## Introduction

Plant physiology and development are prominently affected by changes in ambient temperatures. With the current global climate change, temperatures are gradually shifting and temperature extremes occur more frequently. Predictions of the effect of temperature increment on major crops yield show that each degree-Celsius increase in global mean temperature would cause yield reduction by 3.1-7.4% on average (Zhao et al. 2017). Recent IPCC reports estimate global warming is likely to reach a 1.5°C increase in average surface temperature between 2030 and 2052 if it continues to increase at the current rate, and reach a 2–4^0^C increase by the end of the twenty-first century (IPCC, 2018), thus challenging crop productivity and food security. High temperature is a major abiotic stress that disturbs basic molecular processes, such as protein folding, photosynthesis and assimilates metabolism (Bokszczanin et al. 2013). These effects cause morphological and physiological changes, negatively affecting plant growth and development (Wahid et al. 2007; Bita and Gerats 2013). Yield reduction due to heat stress was documented in various crops such as cereals (wheat, rice, barley, sorghum and maize), pulses (chickpea) and oil yielding crops (mustard, canola) fruits and vegetables (potato, eggplant, cabbage, cauliflower, lettuce, onion, cucumber, musk melon, watermelon and pumpkin) (Hasanuzzaman et al. 2013). When heat stress occurs during the reproductive phase of plant development, the observed consequences include morphological alterations of anthers, style elongation, bud abscission and reduced fruit number, size and seed set. The development of pollen is considered the most heat-sensitive stage (Lohani et al. 2020) as it was shown to be more sensitive than both the sporophyte and female gametophyte tissues (Peet et al. 1998; Young et al. 2004; Wang et al. 2019). Heat stress disrupts of meiotic cell division, abnormal pollen morphology and size, and reduced grain number, viability, and germination capacity (Endo et al. 2009; M. M. Peet et al. 1998; Djanaguiraman et al. 2013; Giorno et al. 2013; Pressman et al. 2002; Firon et al. 2006; Begcy et al. 2019; Prasad et al. 2006). Specifically, pollen viability is considered a central element for heat stress tolerance as high temperatures were shown to impair pollen viability in numerous crop species such as wheat (Begcy et al. 2018), rice (Jagadish et al. 2007), sorghum (Djanaguiraman et al. 2018), soybean (Djanaguiraman et al. 2013), and tomato (Firon et al. 2006), leading to male sterility and reduced fruit/grain production.

Tomato (*Solanum lycopersicum*), an important vegetable crop worldwide, cultivated in a wide range of agro-climatic regions, is very sensitive to heat stress. The tomato fruit set is optimal when the average day and night temperatures range between 21°C - 29°C and 18°C - 21°C, respectively (Pelzer 2008). Prolonged stress of day temperatures exceeding 32°C with night temperature above 20°C cause reduced fruit set, fruit weight, total yield and seed production (El Ahmadi and Stevens 1979; Peet et al. 1998, Sato 2000; Firon 2006). In tomato, pollen heat stress related damage, exhibited by morphological alterations and reduced pollen viability and germination rates, was observed after short episodes of high temperatures at 40°C, or after chronic exposure to milder heat stress of 31-32°C/25-28°C day/night for several months (Firon et al. 2006; Iwahori 1966; Giorno et al. 2013). The decrease in pollen viability and/or germination was shown to cause a significant decrease in fruit set (Iwahori 1965; Rudich et al. 1977; Abdul-Baki 1992; Sato et al. 2000), therefore pollen viability was used as a screening approach to identify heat stress tolerant tomato genotypes. Consequently, several tomato genotypes were identified, that maintain a higher level of pollen viability under heat stress conditions (Dane et al. 1991; Paupière et al. 2017; Driedonks et al. 2018). Pollen viability is therefore often used as a measure of thermotolerance, establishing the correlation between pollen viability and fruit (Pressman et al. 2002; Xu et al. 2017; Pham et al. 2020; Rutley et al. 2021; Firon et al. 2006).

In contrast to the wealth of data demonstrating the correlation between pollen heat stress damage and fruit set, examples of heat stress tolerance/sensitivity not correlated with pollen viability are very scarce. To the best of our knowledge, only two such cases were described. Gonzalo *et al*. (2020) performed a population screen of introgression lines from the wild species *Solanum pimpinellifolium* for reproductive traits under controlled heat stress conditions, and no correlation was found between pollen viability and fruit set (Gonzalo et al. 2020). In a more recent study, Ayenan et. al (2021) screened a collection of 42 cultivated and wild tomato genotypes with good yield components under long term mild heat stress and did not find association between the proportion of viable pollen and fruit set percentage (Ayenan et al., 2021). In this paper, we present yet another example for heat stress tolerance that is not correlated with pollen viability, in a processing cultivar of tomato.

Tomato processing cultivars are used by the food industry to produce tomato paste and sauces, canned crushed, diced, or peeled tomatoes and various juices and soups. For these purposes, breeding companies developed cultivars suited for mechanical harvesting and canning processes. These cultivars are characterized by a determinate growth habit, synchronized fruit set and firm flesh (Hanna 1971; Gould et al. 1992), unlike the indeterminate fresh market cultivars, grown primarily in greenhouses or other covered facilities. Processing tomato plants are cultivated only in open fields, where heat stress conditions are prevalent. Particularly in the Mediterranean basin, including the major tomato producers Italy and Spain, the growing season starts in March–April, when the probability of high temperatures during the sensitive reproductive stage is very high (http://www.wptc.to). However, information regarding the response of processing cultivars to heat stress is very limited.

Here, we characterized the heat stress response of two processing tomato cultivars, which are usually grown in open field conditions therefore exposed to a combination of stress factors, including heat stress, during the reproductive stage. We show that the constant difference in yield between these cultivars is attributed to high temperature conditions. In order to gain information specifically for the response to heat-stress, the same cultivars were tested in a controlled greenhouse, under heat stress and control conditions in a parallel setup. This setup allows the identification of specific heat stress related traits, which is not possible under the uncontrolled, multi-stress field conditions. Our results demonstrate a clear difference in performance under heat stress, which is, unexpectedly, not related to pollen viability.

## Materials and methods

### Plant material and growth conditions

Two tomato (*Solanum lycopersicum*) commercial processing cultivars H4107 and H9780 (Green Seeds Ltd.), were grown during 2018 in three different experimental fields, in different locations as follows: 1. ‘Upper Galilee’ site, at the Northern part of Israel (33°10′50.6″N latitude 35°34′49.6″E longitude; Field size 100 plants), 2. ‘Eden’ site (32°27′58.2″N latitude 35°29′12.2″E longitude; Field size 80 plants) and 3. ‘Volcani’ site at a central region of Israel (31°59′34.6″N latitude 34°49′01.8″E longitude; field size 40 plants). The two cultivars were grown in a completely randomized design in 3-5 replicas (plots). Seeds were sown in germination trays and transplanted in open fields after three weeks. Mature plants were maintained under standard horticultural practices. During the whole growing period climatic data were recorded using the weather stations ‘Khavat Eden’, ‘Beit Dagan’ and ‘Mop Tzafon’ located in Eden, Volcani and Upper Galilee fields, respectively. In addition, the two cultivars were grown in climate controlled greenhouses at the Naan site of Evogene LTD company. In this controlled experiment, four plants from each cultivar were grown under moderate chronic heat stress (MCHS) conditions (32°C-22°C day-night, starting at flowering) and control conditions (25°C-18°C day-night), in a randomized setup, identical between the two rooms. The seeds were sown in germination trays and transplanted into 10L pots filled with soil 21 days after sowing.

### Reproductive traits evaluation

Fruit set and fruit production were evaluated in all three experimental fields and in the controlled experiment. Fruit production (FW – fruit weight) was evaluated by weighing total red-ripe fruits per repeat (plot or plant in the field or controlled experiments, respectively). Fruit set ratio (FS) was evaluated from 10 randomly selected inflorescences from each plot in the field experiments. In the controlled experiment, FS was evaluated from three randomly selected inflorescences in 4 different plants (a total of 12 inflorescences per cultivar). Seed number per fruit (SN) was examined by seeds extraction using three fruits from five plants (Volcani field) or three fruits from five plots (Upper Galilee field). In the controlled experiment, 5-25 fruits from all four plants were sampled. Seeds were extracted using the sulfuric acid method; the locular gel containing the seeds was extracted and soaked in 2% sulfuric acid solution. After 3 hours, the seeds were transferred into a net bag and rinsed under tap water. Seeds were then thoroughly dried in the open air for few days. Seed number was calculated using the weighing method: a small portion was manually counted and weighed, and then the total amount of seeds was estimated by weighing.

### Pollen viability analysis

For pollen viability analysis, flowers at anthesis were collected in the morning (7 to 10 am). In total, three flowers per plant were collected and three plants were used per cultivar. Each anther was cut into two pieces and put in a 1.5 mL tube filled with 0.5mL germination solution [1 mM KNO_3_, 3 mM Ca (NO_3_)_2_·4H_2_O, 0.8 mM MgSO_4_·7 H_2_O, 1.6 mM H_3_BO_3_; (Pressman et al. 2002)], followed by 20 μl of Alexander dye. The Alexander dye consisted of 20 ml of ethanol, 20 mg of malachite green, 50 ml of distilled water, 40 ml of glycerol, 100mg of Acid fuchsin, 2 gr Phenol, and 2 ml of Lactic acid for a 100 ml solution (Alexander 1980). Samples were observed under Leica DMLB epi-fluorescence microscope (Germany) using BF filter, magnified by 10-20. Three fields containing representative pollen pattern were captured with DS-Fi1 digital camera using NIS-Elements BR3.0 software (Nikon). Viable (purple) and non-viable (blue-green) pollen grains were counted manually in ImageJ version 1.43 software using the ‘Cell counter’ plugin (Schneider et al. 2012).

### Statistical analysis

One-way ANOVA was employed to identify significant differences (p<0.05) between the cultivars for each trait. When ANOVA identified significant differences among genotypes, we used the student t-test method as an exact test for all differences between means. These conservative procedures limited the probability of rejecting a true null hypothesis to the desired (p<0.05) level. All statistical analyses were performed using JMP Version 3.2.2 (SAS Institute, Inc., Cary, N.C.).

## Results

### Consistent difference in yield between H4107 and H9780 across multiple years and locations

Following a survey of processing tomato field-testing data from 15 years (between 2005 and 2019) across 17 different locations (Table S1), we detected a consistent difference between two cultivars, i.e., H4107 and H9780. While the yield of H4107 was always above the test average, the yield of H9780 was always lower than the test average (Table S1). When we compared the results of specific years and locations where both cultivars were tested simultaneously, the average yield was 12.4 and 10.8 k/m^2^ for H4107 and H9780, respectively, providing a significant difference (Figure 1a, b). We aimed to understand the source of this difference in order to promote breeding efforts for high yield in field-grown processing tomato. Since the field environment imposes various stresses to the plants, and tomato being particularly sensitive to elevated temperatures, we set to test the possibility that the high temperature conditions usually prevalent in those regions are causing the difference in yield.

**Figure 1.**
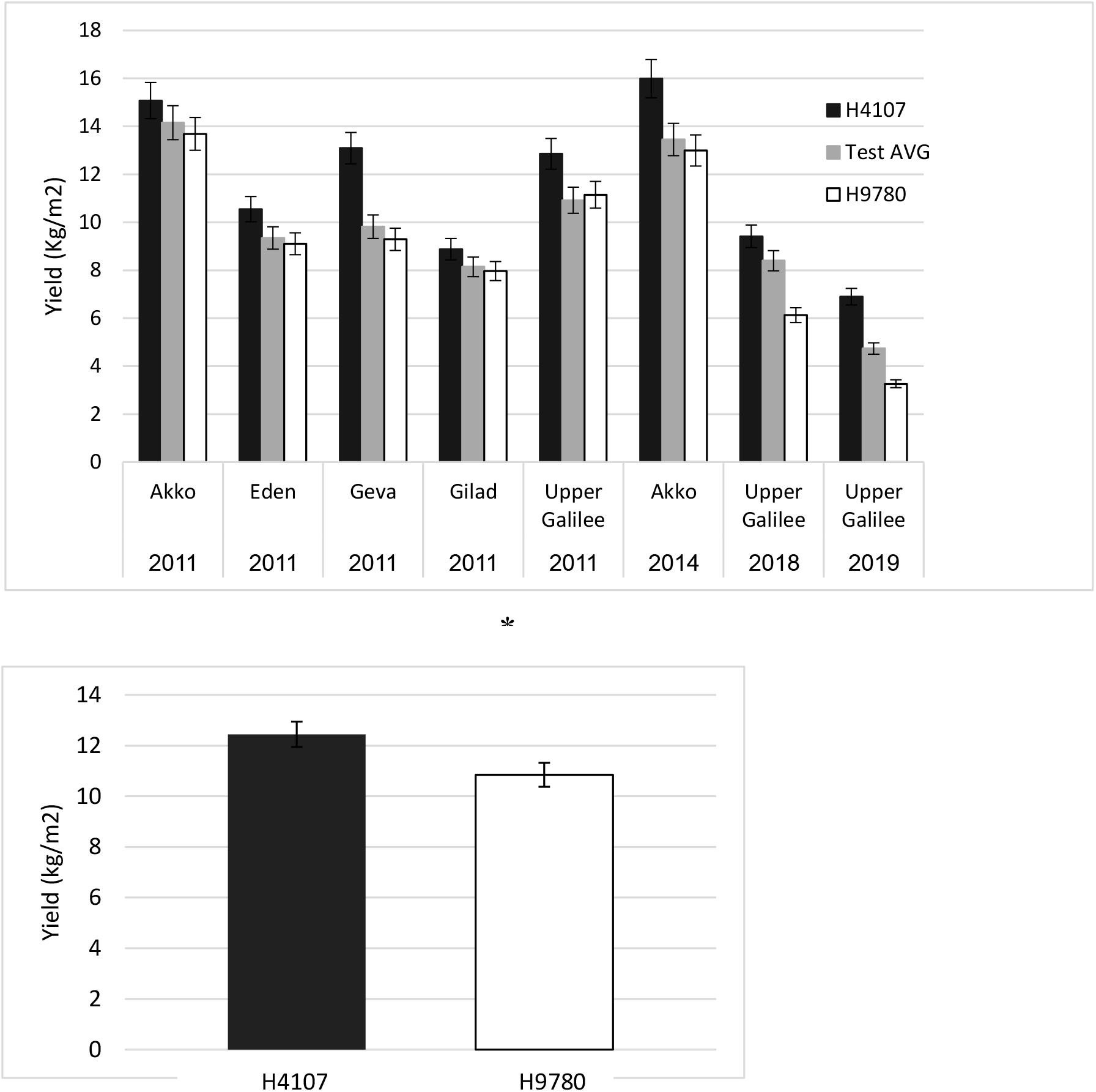
Consistent difference in yield between H4107 and H9780 across years and locations. (A) Average yield of H4107 and H9780 in years and locations testing both cultivars. The test average obtained by yield measurements of multiple cultivars is presented as well. (B) Average yield of H4107 and H9780 across years and locations presented in A. *, statistically significant difference (P-value < 0.05).

### The difference in yield between H4107 and H9780 is associated with high temperature conditions

To test whether the observed difference in yield between H4107 and H9780 is due to their differential response to high temperature, we set field experiments in two locations that are routinely used for processing tomato cultivation, however, differing by their environmental conditions. The ‘Upper Galilee’ field is located in a region that is characterized by hot days and cooler nights during the processing tomato season (May-July), whereas the “Eden” field is located in the Jordan Valley which is characterized by high day and night temperatures, and high humidity. For this reason, planting in “Eden” starts earlier (February until May), to avoid extreme heat stress and yield losses. In addition, we set a small experimental field at the Volcani Center, located in a more temperate region. Overall, we tested the plants under field conditions in three different environments. Environmental data were obtained for each field from a local meteorological station, enabling recording temperature every 3 hours, hence we calculated day and night average and maximum temperatures. Considering that tomato plants experience heat stress when day temperature exceeds 32°C and night temperature exceeds 20°C, our analysis shows that heat stress conditions were indeed prevalent in all three locations, though with some differences (Figure 2a-d). In the Eden field, due to the early planting, heat stress conditions developed around 50 days after flowering. Nonetheless, day and night maximal temperatures surpassed threshold values already 5 days after flowering, generating heat stress conditions throughout the entire reproductive period. In the Upper Galilee field, daily average temperatures were around 32°C, reaching a maximum of approximately 35°C in most days, including three incidences of above 40°C. Night temperatures in the Upper Galilee field were higher than 20°C throughout the period, reaching a maximum of over 30°C on several occasions, presenting more severe heat stress than in the Eden field. Lower temperatures were observed in the Volcani field, where the daily average was usually under 32°C, with four exceptional heat waves. Night temperatures were still high averaging around 25 °C throughout the tested period, thus the plants in the Volcani field also experienced heat stress conditions (Figure 2a-d). Under the above-described conditions, we found that the yield of H4107 was significantly higher than that of H9780 in all fields (Figure 2e), in agreement with our analysis of multiple years and locations data (Figure 1). While H4107 produced 9.0, 6.9, and 11.0Kg fruit/m^2^ in Upper Galilee, Volcani, and Eden, respectively, H9780 produced 5.1, 3.3, and 8.0Kg fruit/m^2^ in the same respective fields. Moreover, yield levels in both cultivars were higher in Eden than in the Upper Galilee and Volcani fields that experienced a more substantial heat stress, suggesting that yield levels are indeed affected by the high temperatures in these locations. The reproductive difference between H4107 and H9780 was further demonstrated by testing fruit set ratio and seed production in the Upper Galilee and Volcani fields (Figure 3). In these locations, H4107 reached 28% and 35% fruit set, respectively, while H9780 had 17% fruit set in both locations (Figure 3a). Similarly, H4107 produced a higher number of seeds per fruit versus H9780, reaching 244 and 96, respectively, in the Upper Galilee field. In the Volcani field, H4107 had on average 61 seeds per fruit, and H9780 produced only 21 seeds per fruit on average, maintaining a significant difference (Figure 3b).

**Figure 2.**
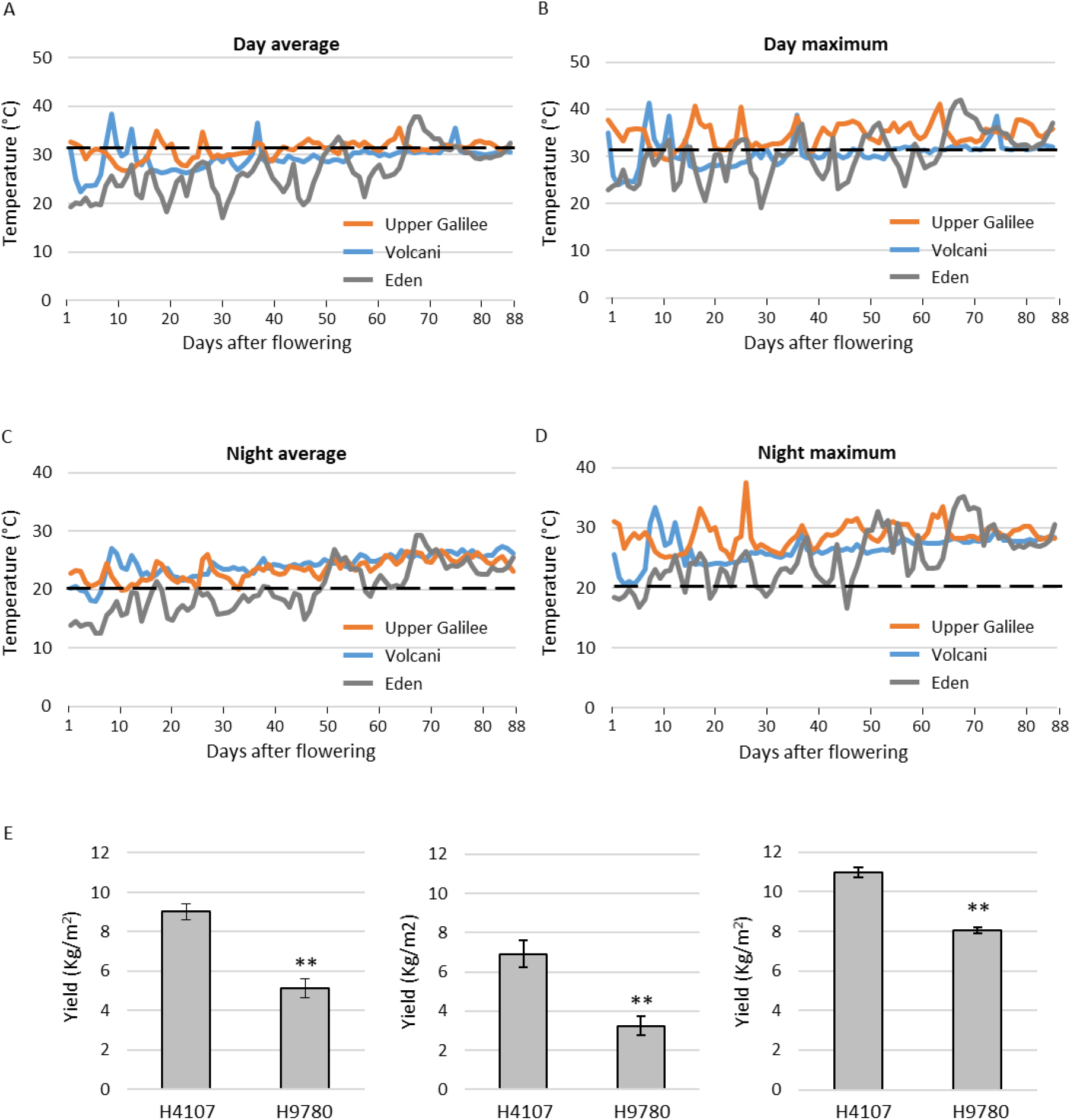
Field experiments conditions and yield. Temperatures were recorded constantly in the three experimental sites: Upper Galilee, Volcani and Eden. Daily average (A), daily maximum (B), night average (C) and night maximum (D) were calculated for the reproductive period and are presented from the first day of flowering until the end of the experiment (88 days after flowering). (E) Yield performance for H4107 and H9780 in the Volcani (left), Upper Galilee (middle) and Eden (right) fields. **, statistically significant difference (P-value < 0.01).

**Figure 3.**
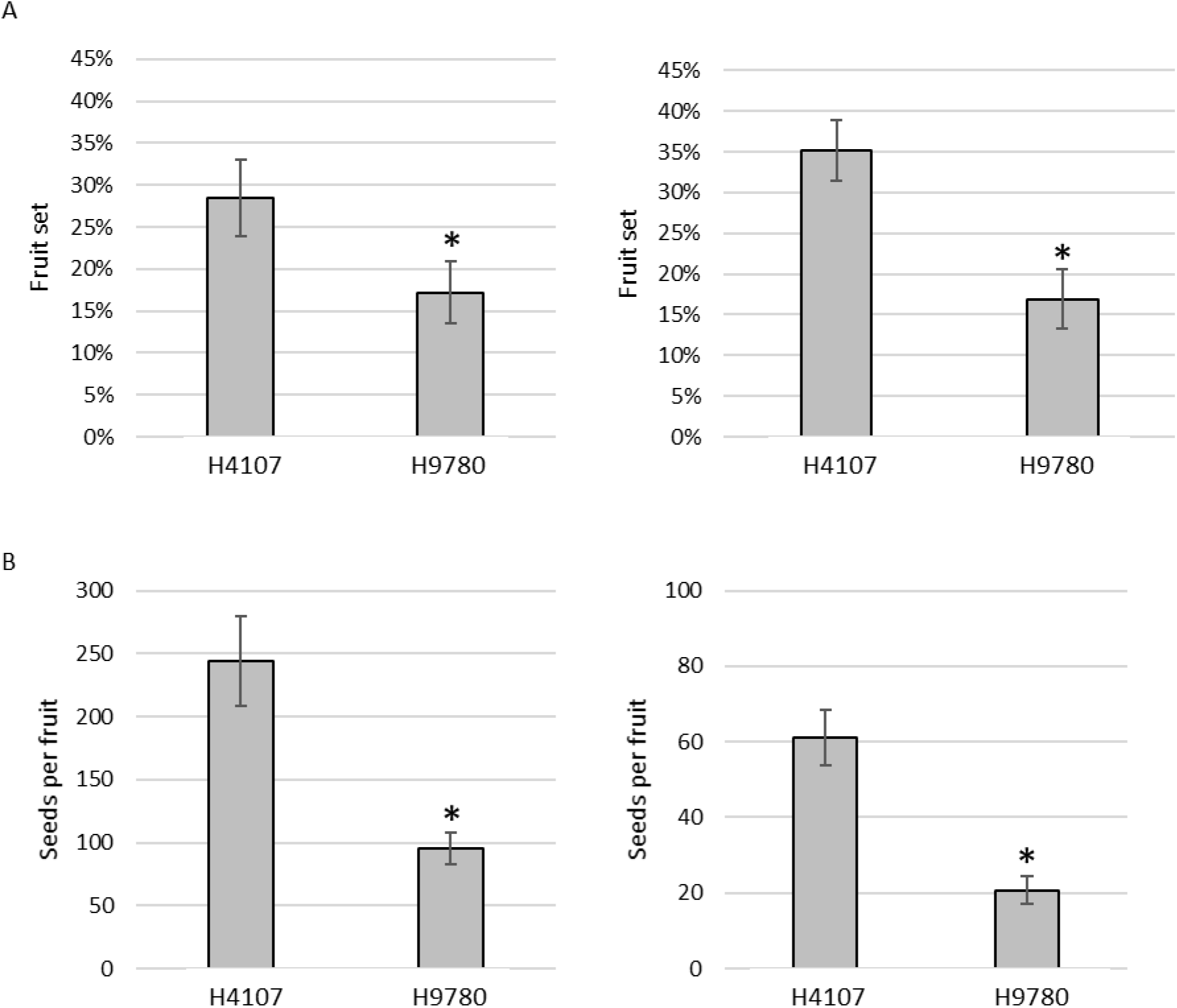
Fruit set and seed number measurements in the field experiments. (A) Fruit set rates of H4107 and H9780 in Upper Galilee (left) and Volcani (right) fields. (B) Seeds number per fruit for H4107 and H9780 in Upper Galilee (left) and Volcani (right) fields. *, statistically significant difference (P-value < 0.05).

In order to validate the effect of heat stress on the productivity of H4107 and H9780, we set a controlled experiment in which the same cultivars were grown under either MCHS (32°C/22°C day/night), or control conditions (25°C/18°C day/night) in separate rooms. At the beginning of the experiment, both rooms were maintained under control conditions. Once plants started to flower, MCHS was initiated in one room while the other room was kept at control conditions throughout the rest of the plants growth (Figure 4a). Fruit set rate and seed production were analyzed under both conditions. We found no significant difference between H4107 and H9780 in both parameters measured (i.e. 64-68% fruit set and 52-92 seeds per fruit) under control conditions. However, under MCHS conditions, H4107 performed better than H9780, as the fruit set was 36% versus 19% in H9780. Seed number per fruit was 71 and 23 for H4107 and H9780, respectively (Figure 4b-c). Markedly, fruit set ratios were very similar between field and controlled heat stress for both cultivars, supporting the occurrence of heat stress conditions in the field experiments. Importantly, these results confirm that the observed difference in yield and other reproductive traits under open field conditions are due to high temperatures, and suggest that H4107 is more tolerant than H9780 to heat stress.

**Figure 4.**
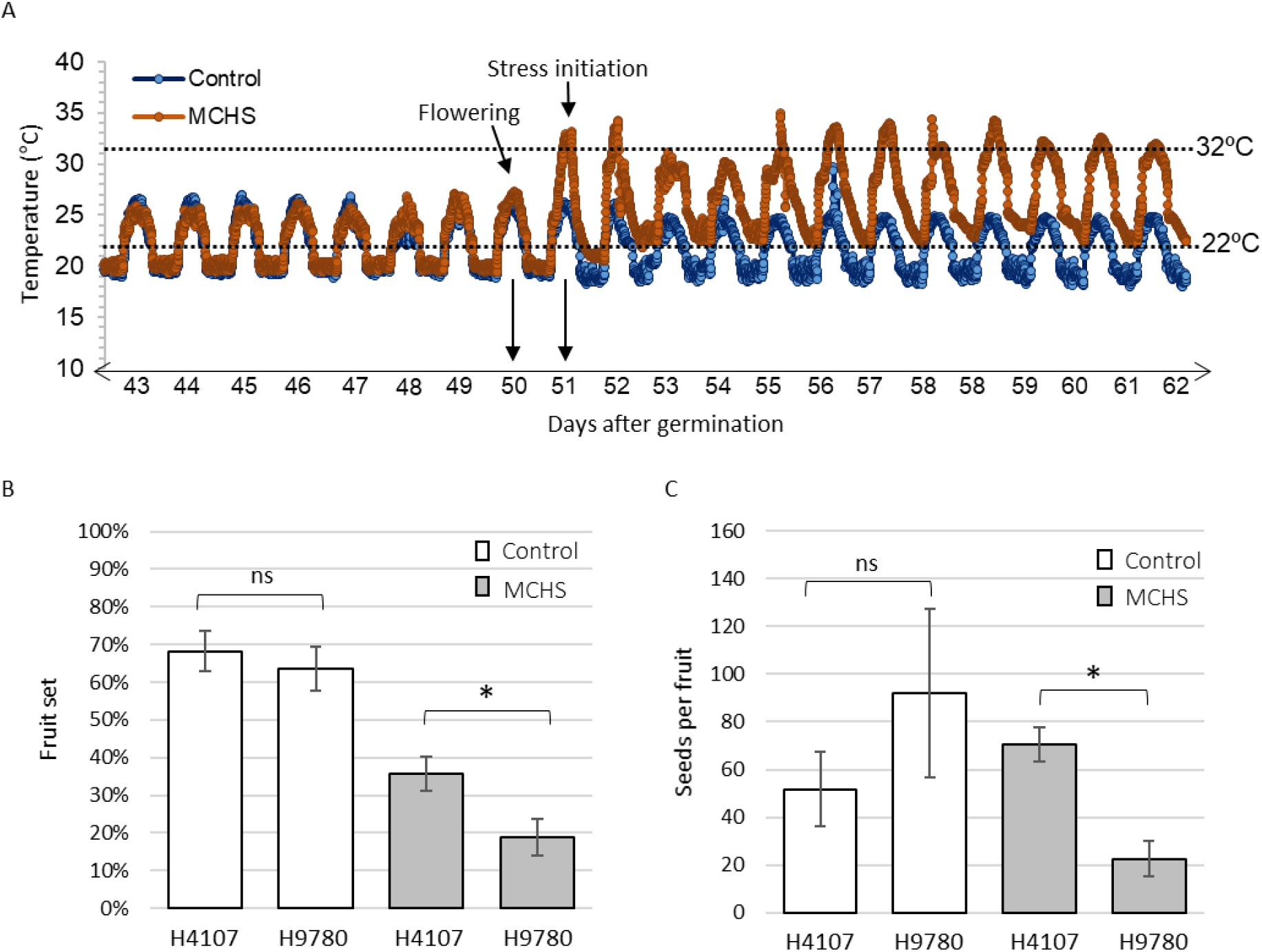
Controlled experiment conditions and reproductive measurements. (A) Temperatures measured every five minutes in both control (blue) and MCHS (brown) greenhouses. Black arrows denote day of flowering and day of stress initiation. Threshold temperatures for heat stress conditions in tomato are marked by dotted lines. (B) Fruit set ratio for H4107 ad H9780 under control (white bars) and MCHS (grey bars) conditions. (C) Seeds number per fruit in H4107 and H9780 under control (white bars) and MCHS (grey bars) conditions. MCHS, moderate chronic heat stress. *, statistically significant difference (P-value < 0.05). ns, not significant.

### The difference in heat tolerance between H4107 and H9780 is not related to pollen viability

Since pollen viability is widely recognized as a main parameter determining plant heat stress tolerance (Dane et al. 1991; Paupière et al. 2017; Driedonks et al. 2018), we aimed to test whether the heat stress tolerance of H4107 can be at least partially explained by higher degree of pollen viability under heat stress conditions. To address that, we analyzed pollen viability percentage in field and controlled conditions. In the Upper Galilee field, we found no significant difference between H4107 and H9780, as both showed 60-70% viable pollen out of total pollen grains (Figure 5a). Pollen viability was lower in the Volcani field (30-45%), yet still similar between the cultivars (Figure 5b). In the controlled experiment, pollen viability reached 90-100%, even under MCHS conditions, and again, similarly between H4107 and H9780. Interestingly, the same levels were found under control conditions (Figure 5c), meaning that pollen viability was not affected by heat stress in these cultivars and is not linked with the heat stress tolerance of H4107. Our results also suggest that the low rates of pollen viability in field conditions is not due to the high temperatures, but rather to another environmental factor.

**Figure 5.**
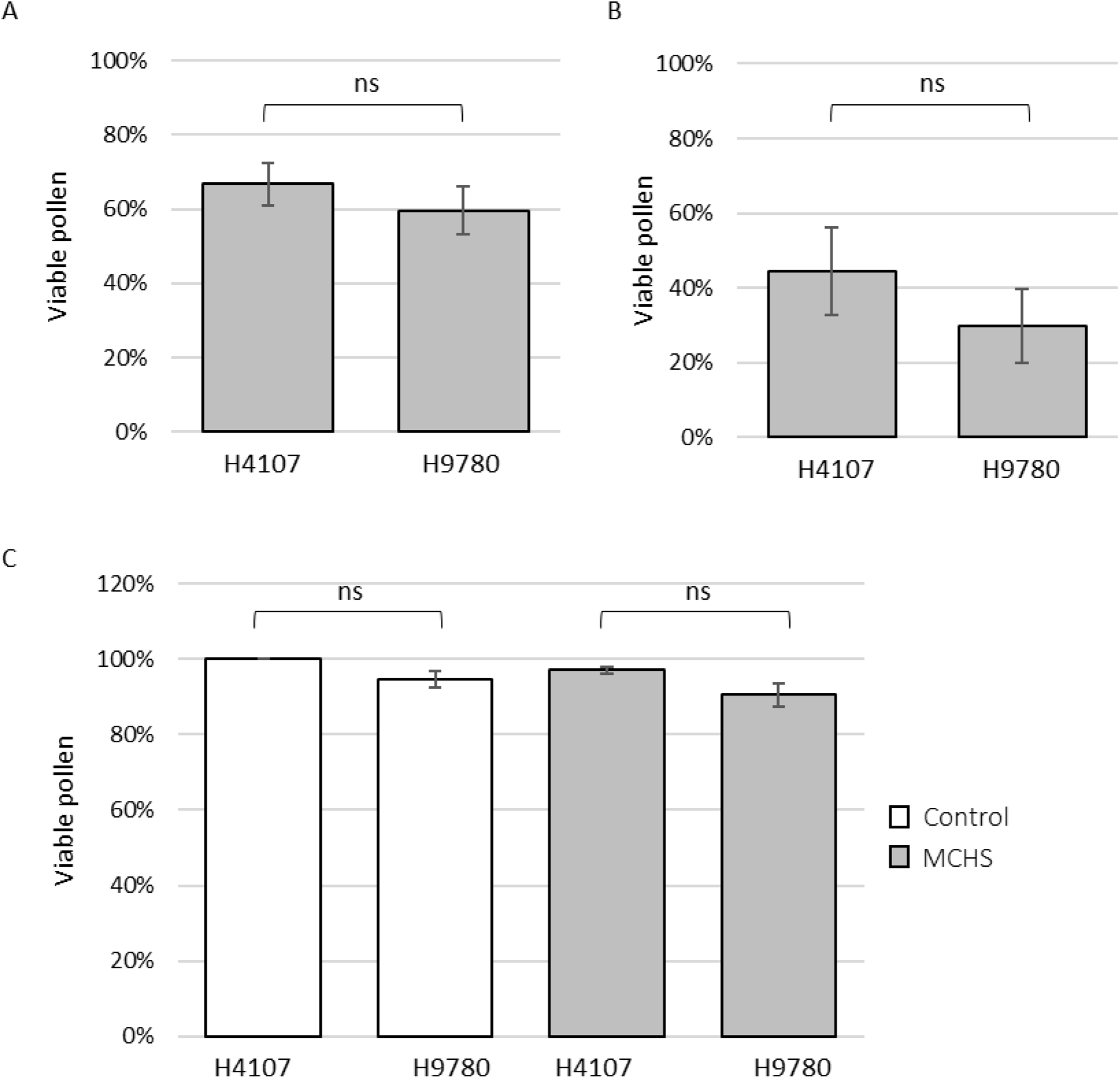
Pollen viability in field and controlled experiments. Percentage of viable pollen from post-anthesis flowers of H4107 and H9780 at the (A) Upper Galilee field, (B) Volcani field and (C) controlled greenhouses, under control (white bars) and MCHS (grey bars) conditions. MCHS, moderate chronic heat stress. ns, not significant.

## Discussion

Current literature on processing tomatoes in general and on their response to heat stress in particular is very limited. We identified a consistent difference in yield between H4107 and H9780 across multiple years and locations. This difference is manifested by higher fruit set rate and total fruit weight of H4107. We found this difference to be associated with the response to heat stress, meaning that H4107 is more heat stress tolerant than H9780, presenting better reproductive performance in terms of fruit set and seed production under high temperature conditions. H4107 was bred and adapted for humid and arid environments by the Heinz company (https://d36rz30b5p7lsd.cloudfront.net/372/studio/assets/v1611911409263_1054604699/2021%20HeinzSeed%20International%20Brochure.pdf), but heat stress tolerance was not reported so far. Interestingly, the heat stress tolerance we observed was not correlated with better pollen viability, implying that other factors mediate the tolerance in this system. In one of the earliest studies on heat stress response in tomato, Levy *et al*. (1978) showed that the characters contributing to low fruit set under heat stress were bud drop and style exertion which were more pronounced in susceptible cultivars. Actually, no fruit set was ever observed when the style protruded out of the antheridial cone (Levy et al. 1978). Fruit setting was correlated with bud abscission and style elongation under field conditions as well (Singh et al. 2015; Kugblenu et al. 2013). Considering this aspect, we tested bud abscission and style elongation ratios in field and greenhouse but no significant difference was found between H4107 and H9780 (data not shown). Alternatively, ovule development and post-pollination interactions were also demonstrated to negatively influence fruit set, by applying pollen from control condition flowers onto freshly open flowers grown under heat stress conditions (Peet et al., 1997; Xu et al., 2017).

Ayenan et. al (2021) showed recently that in some tomato genotypes grown in the greenhouse, pollen viability was not correlated with fruit set and yield (Ayenan et al., 2021). On the same hand, our results suggest that while pollen viability is a valid trait demonstrating heat stress tolerance in various tomato genotypes, it may not be the case in open field processing cultivars. If this is due to their genetic structure or the complex environment they were bred in, or a combination of both, is yet to be determined following a comprehensive follow-up study.

Generally, in plant science research, field and greenhouse data are inconsistent, explained by the big difference in environmental conditions between the two experimental systems. We found that fruit set is highly similar between the controlled experiment (36% and 19% for H4107 and H9780, respectively) and the field experiments (28-36% and 17% for H4107 and H9780, respectively). Thus, our results demonstrate consistency in regard to a complex trait (yield), suggesting that in our system, controlled greenhouse experiments are highly relevant for agricultural conditions, facilitating translating research from lab to practice. Moreover, our results demonstrate the importance of temperature-controlled experimental systems in isolating specific heat-stress related phenomena.

In order to address the challenge of maintaining crop productivity in areas of temperature increment, the development of thermo-tolerant cultivars is needed. To achieve that, a comprehensive understanding of the agronomical, physiological and molecular responses of crop plants to heat stress is vital (Berry and Bjorkman 1980; Brestic et al. 2018). In light of the research presented here, which demonstrates a unique feature of specific cultivars, emphasis should be put on local and relevant cultivars that may offer different attributes in terms of response to the environment.

## Acknowledgements

(removed)

Authors declare no conflict of interests.

---

**Table S1.**
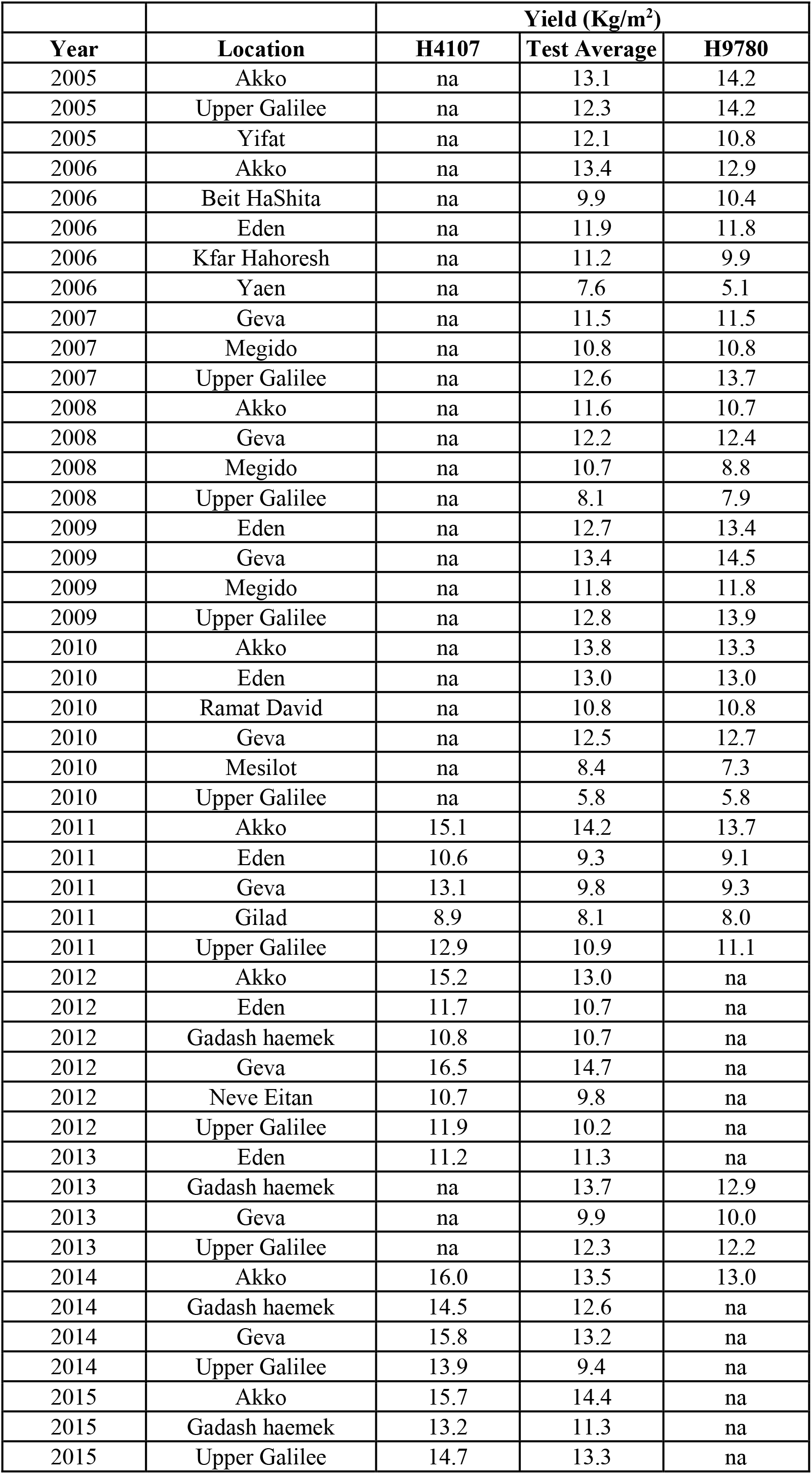
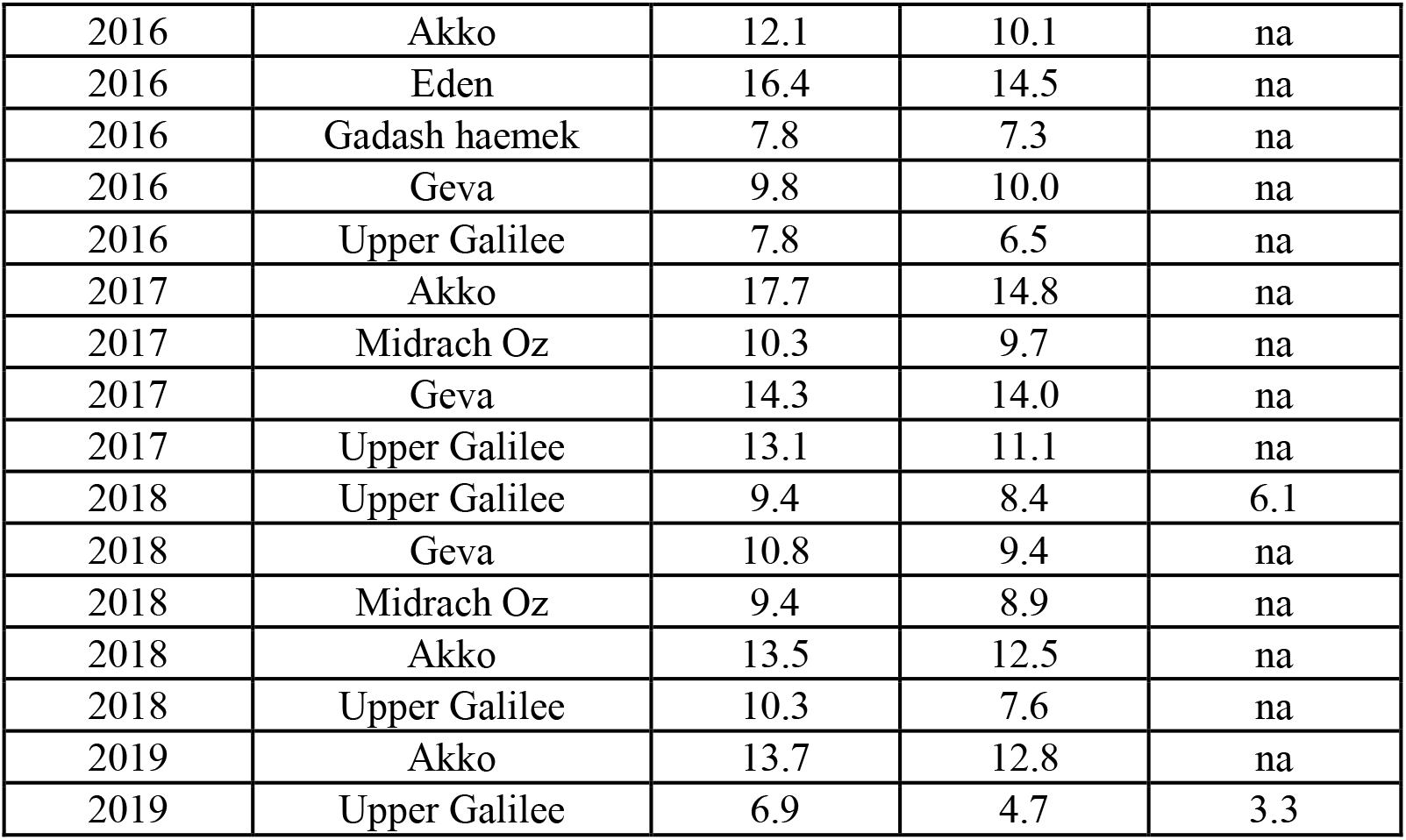
Yield measurements in field trials of processing tomatoes between 2005 and 2019 in different locations. Presented here are yield values for H4107 and H9780 as well as the whole test average. na, not applicable – the cultivar was not tested

